# The Effect of post Chlamydia Trachomatis Infection treatment on Reactive Oxygen Species and Sperm Parameters of Infertile Men

**DOI:** 10.1101/2020.10.21.349811

**Authors:** Reza Azmakan, Reza Salman Yazdi, Faramarz Farahi, Vahid Esmaeili, Seyed Kazem Bidoki, Saied Jafari

## Abstract

**Background:** Chlamydia trachomatis (CT) infection is often mentioned as a silent disease. Reactive oxygen species (ROS) can also cause Sperm apoptosis and have negative impact on Sperm parameters. The objectives of this study were to elucidate the association between Sperm parameters and ROS caused by CT infections resulting in male Infertility as well as evaluating the role of antibiotic therapy.

**Materials and methods:** A total of 848 infertile males having normal and abnormal Sperm parameters were included. After Semen sampling, the CT IgA were measured by Elisa and confirmed by Nested PCR. ROS was determined by Chemiluminescence. After treatment under the direct supervision of the private urologists. Then, the second Semen samples were taken and subjected to tests on Sperm parameters and ROS levels as assessed again.

**Results:** The levels of ROS and morphology were improved following the treatments (P<0.05). Antibiotic therapy due to CT infection, could reduce ROS, improve normal morphology and recover some of Semen parameters.

**Conclusions:** Our findings indicate that CT infection and Sperm parameters were associated with the rate of ROS in infertile men. However, after treatment, ROS value dropped allowing the recovery of certain Sperm parameters. Antibiotic therapy can improve some Semen quality parameters and treat the male Infertility.

Reza Azmakan, Department of Andrology, Reproductive Biomedicine Research Center, Royan Institute for Reproductive Biomedicine, ACECR, Tehran, Iran, https://orcid.org/0000-0001-6718-3348.

## BACKGROUND

It has been reported that about 10 to 15 % of couples suffer from Infertility due to different incentives. Bacterial infections of the reproductive system are one of the causes of Infertility (8 to 35 %) (Oehninger, 2000). Infection could lead to deterioration of Spermatogenesis, impairment of Sperm function and/or obstruction of the seminal tract. The major pathogens are Chlamydia trachomatis, Neisseria gonorrhoeae, and some genital Mycoplasma (De Francesco et al., 2011). Chlamydia trachomatis causes at least 50% of common bacterial Sexually transmitted infections in the world (Stamm et al., 2007). Infection of Semen and male sex gland with CT is associated with was increased ROS levels, was increased motility and Sperm morphology which are important factors in men Infertility. Chlamydia trachomatis is known as a silent disease. Reactive oxygen species (ROS) including oxygen ions, free radicals and peroxide is produced by Sperm and leukocytes in the Semen (Agarwal et al., 2006). ROS has a dual role in male fertility. On one hand carrying capacity, acrosome reaction and fertilizing, and on the other hand causing severe damages to the Sperm. Effects of Infertility caused by ROS are; Sperm membrane damage, reduced Sperm mobility and inability to fuse with the oocyte cell. ROS can also alter Sperm DNA, pass through the damaged DNA paternity and reach embryo. Was increased ROS can result in disruption of the structure and increase permeability that the lack of fertilization and abnormally shaped cells will follow (Tirado et al., 2003). Chlamydial IgA antibodies in seminal fluid can activate the immune system and the ROS leads to was increased lipid peroxidation of Sperm membrane and Spermatozoa apoptotic effects (Mahfouz et al., 2010). Reduced Sperm motility, is caused by H_2_O_2_ penetration into the cell and inhibits the activity of some enzymes, which leads to was decreased phosphorylation of axonemal protein and was decreased Sperm motility (Schillinger et al., 2005).

Sensitivity and specificity of molecular methods, especially Nucleic Acid Amplification Test (NAAT) is high for the diagnosis of chlamydial infection in Semen inside urine (de Barbeyrac et al., 2006). Diagnosis of chlamydial infection and methods for measuring ROS are done in a variety of ways. Methods of measuring ROS such as; Direct method-measurement of MDA by thiobarbituric LPO (Tavilani et al., 2005), Measuring isoprostanes (Khosrowbeygi & Zarghami, 2007), Urinary 8 OHDG (Bohring & Krause, 2003), Sperm chromatin dispersion test (SCD) (Muriel et al., 2006; Organization, 2010), COMET method, TUNEL as well as indirect methods; Chemiluminescence methods-measuring superoxide ions produced by lucigenin sensitive probes which is an indicator to quantify the oxidation activity.

The aims of this study were to compare the frequency of genital C. trachomatis (with normal and abnormal Semen parameters), to elucidate the association of asymptomatic infections with male Infertility, to evaluate the effect of antibiotic treatment on improvement of Spermatozoa parameters, and to examine ROS levels and Sperm parameters before and after treatment in infected infertile men.

## RESULTS

### Study population

The primary sample size was 848 Semen samples. Samples were taken from men with normal and abnormal Semen (low Sperm count, leukocytoSpermia, low Sperm progressive motility, low normal morphology). Of the 848 input samples, 10 samples tested by Elisa positive C. Trachomatis were reported. Eight positive samples (by nested PCR) were certain that one sample was excluded due to non-recourse and 7 samples were analyzed. Sperm parameter and ROS level were compared before and after treatment with antibiotics (7 patients). The average age of peoples in this study was 31.8 ± 0.5 years (ranging from 21 to 57). The mean duration of Infertility for study group was 58.2 ± 2.9 month.

### Frequency of C. trachomatis between the study groups

Of 848 patients (with normal and abnormal Sperm) included in the study group (infertile men), seven (0.83%) were tested positive for C. trachomatis. All infected patients in the study group were resolved from the infection after the treatment completion.

### Semen parameters

Table 1, 2 compares the Semen parameters before and after the antibiotic therapy, respectively. Figure 1 shows the Semen parameters in infected infertile men before and after the antibiotic therapy. There was not statistically significant difference in the Semen parameters. The count of leukocytes, Sperm count, total motility, Class A (rapid progressive), class D (non-motile), volume of Semen and pH Semen after treatment, the parameters of recovery showed improvement with no significant differences (P> 0.05.). Motility of the Class B (progressive) and class C (non-progressive) had no significant differences. The mean count of white blood cells in the pre-treatment sample was 0.41±0.518 million per ml, and the mean count of white blood cells in the samples after treatment was 0.09±0.146 million per ml, The count of WBC was decreased after treatment compared with before treatment, but this reduction was not significant (P-value=0.144). The Sperm count was increased numerically after treatment, but this increase was not significantly different (P-value=0.128). The Sperm motility was increased numerically after treatment compared with before treatment, but this increase was not significantly different (P-value=0.398). Sperm motility including class A, class B, class C and class D before and after treatment was not significant (P-value=0.138). The volume of Sperm samples was not statistically significant before and after treatment (P-value = 0.249). The pH Semen samples were not statistically significant before and after treatment (P-value = 0.157). The normal morphology after treatment compared with the baseline was significantly was increased (P-value=0.024).

**Table 1.** Comparison of semen parameters before and after the antibiotic therapy

| Semen parameters | Z-value | P-value |
| --- | --- | --- |
| After ROS total / Before ROS total | 2.023<sup>a</sup> | 0.043* |
| After ROS WBC / Before ROS WBC | 2.023<sup>a</sup> | 0.043* |
| After ROS base / Before ROS base | 2.023<sup>a</sup> | 0.043* |
| After pH / Before pH | 1.414<sup>a</sup> | 0.157<sup>ns</sup> |
| After normal morphology | 2.251<sup>b</sup> | 0.024* |
| After volume / Before volume | 1.153<sup>b</sup> | 0.249<sup>ns</sup> |
| After class D / Before class D | 0.676<sup>a</sup> | 0.499<sup>ns</sup> |
| After class C / Before class C | 0.676<sup>a</sup> | 0.499<sup>ns</sup> |
| After class B / Before class B | 1.014<sup>b</sup> | 0.310<sup>ns</sup> |
| After class A / Before class A | 1.48<sup>b</sup> | 0.138<sup>ns</sup> |
| After sperm motility total / Before sperm motility total | 0.845<sup>b</sup> | 0.398<sup>ns</sup> |
| After sperm count / Before sperm count | 1.521<sup>b</sup> | 0.128<sup>ns</sup> |
| After WBC / Before WBC | 1.461<sup>a</sup> | 0.144<sup>ns</sup> |

a: Based on positive ranks.
b: Based on negative ranks.
c: Wilcoxon Signed Ranks Test
ns: The difference is not significant.
\*: The difference is significant at the 5% level
p < .05 is considered statistically significant. a Study group (infertile men) before antibiotic therapy. ROS, reactive oxygen species; WBCs, white blood cells. P < .05 is considered statistically significant. ROS, reactive oxygen species; WBCs, white blood cells; ROS wbc, ROS base and total ROS levels observed after treatment was significantly lower than before the intervention and after the intervention in normal morphology values; statistically significantly higher than before the intervention (marked with \*). The remaining variables did not show significant difference (with n.s specified).

**Table 2.** Comparison of semen parameters in infected patients of the study group (infertile men) before and after the antibiotic therapy.

| Sperm parameters | Sperm parameters | Before treatment | SEM | After treatment | SEM | P-Value |
| --- | --- | --- | --- | --- | --- | --- |
| WBCs (million/ml) | WBCs | 0.41 | 0.195 | 0.09 | 0.055 | 0.0144 |
| Sperm count (million/ml) | CS | 57.11 | 15.04 | 68.31 | 17 | 0.128 |
| Total sperm motility class A+B+C (%) | TSM | 39.67 | 10.53 | 47.09 | 8.064 | 0.398 |
| Progressive motility class A (%) | PMA | 2 | 0.813 | 7.29 | 3.655 | 0.138 |
| Progressive motility class B (%) | PMB | 20.89 | 5.556 | 22.97 | 6.534 | 0.31 |
| Nonprogressive motility class C (%) | NPM | 16.79 | 5.163 | 15.51 | 3.58 | 0.499 |
| Immotile class D (%) | ID | 60.33 | 10.53 | 54.23 | 8.567 | 0.499 |
| Semen volume (ml) | SV | 3.13 | 0.796 | 3.6 | 0.765 | 0.249 |
| Normal sperm morphology (%) | NSM | 3.86 | 1.243 | 5.14 | 1.317 | 0.024 |
| Semen pH | SP | 7.83 | 0.028 | 7.77 | 0.041 | 0.157 |
| ROS base (RLU/sec/$\times 10^6$ sperm) | ROSB | 62.52 | 6.834 | 30.19 | 1.419 | 0.043 |
| ROS WBCs (RLU/sec/$\times 10^6$ sperm) | ROSW | 82.44 | 10.842 | 33.84 | 1.809 | 0.043 |
| ROS total (RLU/sec/$\times 10^6$ sperm) | ROST | 133.02 | 31.514 | 44.17 | 5.326 | 0.043 |

Data are presented as mean ± standard error of the mean (SEM); $p < .05$ is considered statistically significant. ROS, reactive oxygen species; WBCs, white blood cells.

**Figure 1:**
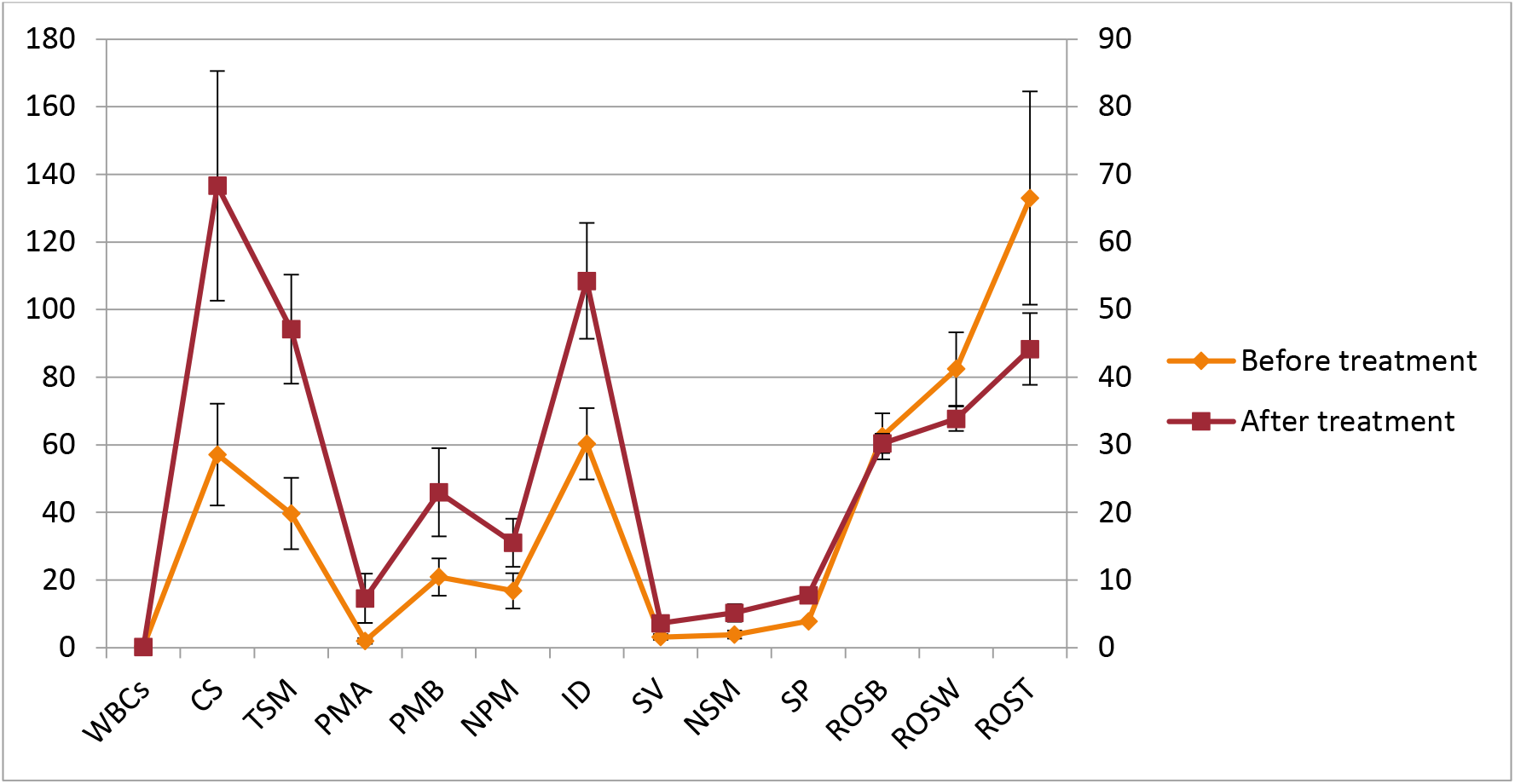
Linear chart for comparison of Semen parameters in infected infertile men, before and after the antibiotic therapy. SV: Semen volume (ml); SC: Sperm count (million per ml); SC: Sperm count (million); TSM: total Sperm motility class A+B+C (%); PMA: progressive motility class A (%); PMB: progressive motility class B (%); TPM: total progressive motility class A+B (%); ID: Immotile class D (%);NPM; nonprogressive motility class C (%); NSM: normal Sperm morphology (%);SP: Semen pH; WBCs: white blood cells (million per ml); ROS B: reactive oxygen species Base(RLU/sec/× 106 Sperm); ROSW: reactive oxygen species WBCs (RLU/sec/× 106 Sperm); ROST: reactive oxygen species total (RLU/sec/× 106 Sperm). Error bars are standard error of the mean (SEM).

### ROS level and Semen WBC

Table 1, 2 compares the ROS before and after the antibiotic therapy and Figure 1 compares the ROS level in infected infertile men before and after the antibiotic therapy. The ROS level was significantly was was decreased after treatment (P-value=0.043). The level of ROS WBCs was significantly was was decreased (P-value=0.043). The total ROS was significantly was decreased after treatment (P-value=0.043). Table 3: no significant correlation was found between Chlamydia infection and white blood cell count (WBC).

**Table 3.** Count of WBC in the samples before and after treatment. In overall comparison, there Leukocyte reduction after treatment, the difference was not significant.

| Mill/ml | Sample 1 | sample2 | sample3 | sample4 | sample5 | sample6 | Sample 7 |
| --- | --- | --- | --- | --- | --- | --- | --- |
| WBC before treatment | 0 | 1 | 0 | 0 | 1 | 0.9 | 0 |
| WBC after treatment | 0 | 0 | 0.3 | 0 | 0.3 | 0 | 0 |

A Values of sperm count per ml as well as total sperm count of only four infected patients in the study group were lower than the WHO normal ranges before antibiotic therapy.

### First and second PCR photo from gel agarose using primers F1-R1 and F2-R2

The results (Figure 2 and 3) Showed that ROS base and ROS, WBC and total ROS and Normal morphology were significantly improved after treatment (P<0.05). Variables such as; count of leukocytes, Sperm count, motility (total), Class A (rapid progressive), class D (immotile), volume of Semen, and Semen pH showed improvement but this improvement was not statistically significant (P> 0.05.). In variables such as motility of the Class B (progressive) and class C (non-progressive) no differences were observed.

**Figure 2:**
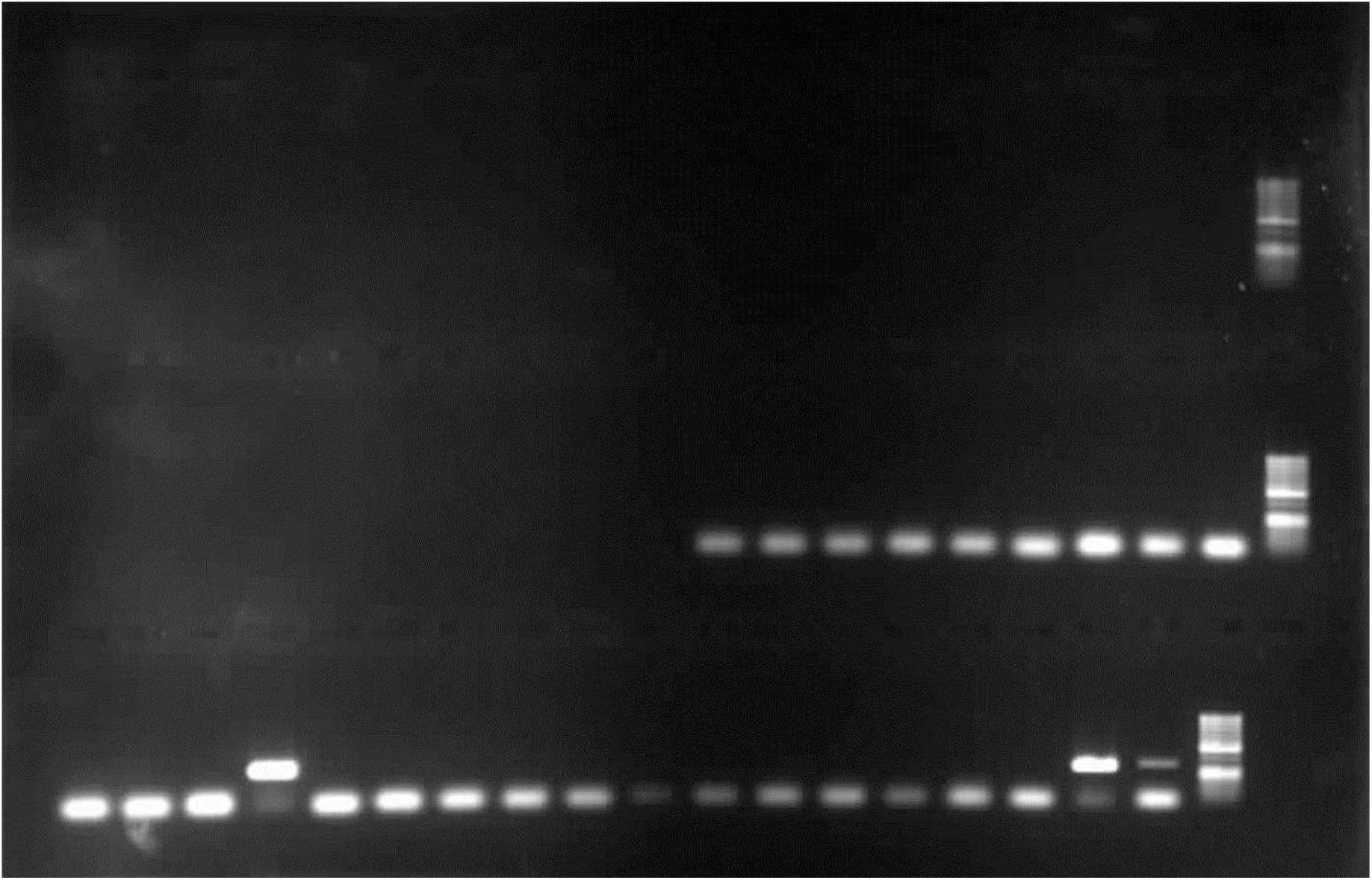
The second PCR 1.0 % gel agarose using primers F2-R2 (317 bands bp) First row from left to right as follows: Chlamydia trachomatis DNA Ladder 50bp-positive control (317 band bp)-positive samples (317 band bp) −12 sample negative - positive samples (317 band bp) −3 negative samples 2nd row from left to right as follows: DNA Ladder50bp - 9 negative samples

**Figure 3:**
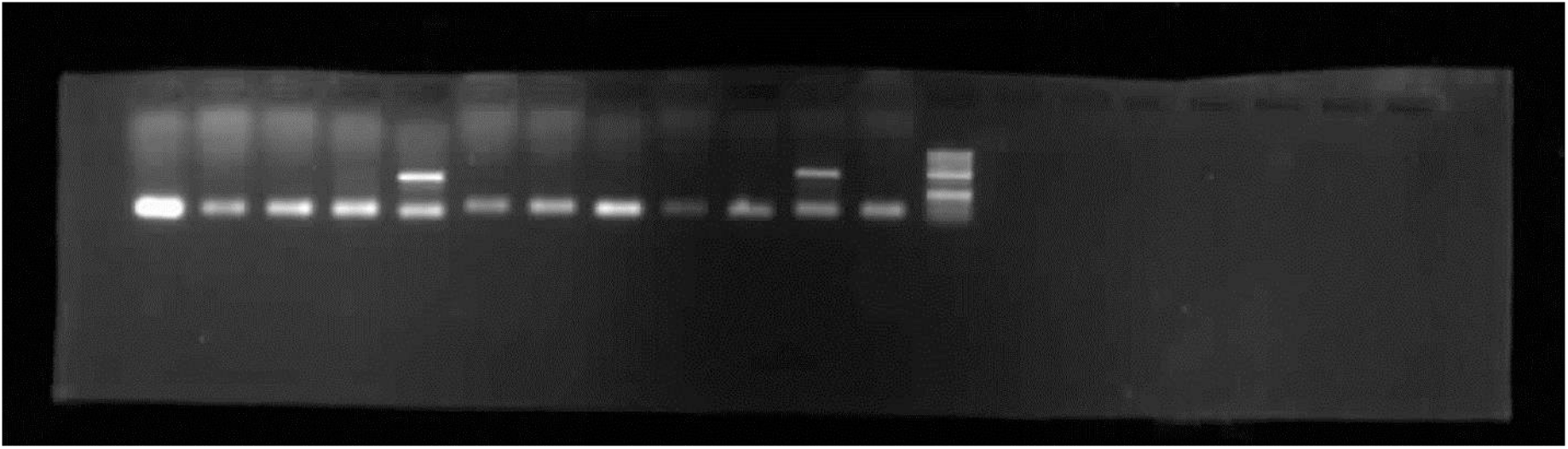
The first PCR photo from 1.0 % gel agarose using primers F1-R1 (517 band bp) From left to right are as follows: DNA Ladder 50bp - positive sample Negative sample (517 band bp) - 5 samples positive sample Negative (517 band bp) - 4 negative samples

## Conclusion

In conclusion, medication with antibiotics showed to be effective in infected patients whose Nested PCR was positive. The quality of natural Sperm morphology and some other Semen parameters improved. ROS level was also was decreased. The results of the present study indicated that asymptomatic infections caused by CT were associated with male Infertility, and high WBC and abnormal Sperm parameters do not indicate infection with this bacterium. Our findings indicate the importance of screening programs for asymptomatic infections due to this bacterium and its treatment. Our findings will also be helpful for infected patients who have abnormal Semen parameters to maintain fertility and reproductive health.

## Discussion

The results showed that chlamydia infection and the rate of ROS and Sperm parameters were co-related in the infertile men (those tested PCR positive), because after medication and infection (negative PCR test after treatment) the rate of ROS was decreased and some Sperm parameters had been improved. Chlamydia is most likely a direct adhesion to the Sperm causing Infertility (Al-Mously et al., 2009). In 50-80% of cases are asymptomatic infection (Hosseinzadeh et al., 2004). Anyone with Chlamydia trachomatis infection should be followed up 3-12 month of treatment (Aghaizu et al., 2014). Chlamydia trachomatis causes common bacterial STD in the world and responsible for at least 50 % of the causes of pelvic inflammatory disease and 87 % of men and women infection. Chlamydia trachomatis is the most common pathogen identified in the nongonococcal urethritis (NGU) group, with up to 42% of NGU cases being caused by this bacterium (Stamm et al., 2007). There is an uncertainty about the role of C. trachomatis infection in Infertility and free radicals of oxygen (ROS) and even the type of sample (urine, Semen) as well as the type of diagnostic method. It was reported that increasing the count of leukocytes in Semen was correlated with poor Semen quality (Kalayoglu & Byrne, 2006). In men with chlamydial IgA positive, high percentage of dead Sperm and seminal fluid was observed in the prevalence of leukocytes (Idahl et al., 2007). In our study, although there was a relationship between the count of leukocytes with chlamydial infection, however, this relationship was not statistically proven. Although the method of NAAT for detection of genital C.trachomatis in women’s and men’s have been reported (Hislop et al., 2010) but the PCR test were more sensitive, which could be used as a confirmatory test for C. trachomatis infection. Combination of different methods screening for C. trachomatis should be used in research because there is no approved method for Chlamydia in Semen (Peeling & Embree, 2005). PCR and ligase chain reaction are more sensitive (Watson et al., 2002). The diagnostic method used in our study was close to the studies conducted by Hosseinzadeh, Embree and Templeton, in which, the first was by Elisa and confirmed by Nested PCR. In the study of Hosseinzadeh two primer pairs were used for the diagnosis of C. trachomatis and our study was consistent with these studies (Hosseinzadeh et al., 2004). Our results were also consistent with the study conducted by Elisa and PCR (Kalantar et al., 2007). C. trachomatis in prostatitis patients affected parameters such as Sperm concentration, percentage of Sperm motility (Mazzoli et al., 2010). Men with CT infection showed poor Semen quality (Al-Mously et al., 2009) (Hosseinzadeh et al., 2004)

The association between CT infection and Semen weakening was observed (Cunningham & Beagley, 2008) (Bezold et al., 2007) (Gallegos et al., 2008) (Ahmadi et al., 2018) (Shiva et al., 2011). Some studies suggested that some bacterial species had negative effects on Sperm parameters including motility and morphology (Berktas et al., 2008) (De Francesco et al., 2011) (Prabha et al., 2010) (Mazzoli et al., 2010; Zhang et al., 2011) (Eley et al., 2005; Mazzoli et al., 2010). Those results were consistent with the results of us in the current study. Two studies by Motrich showed that CT infection altered Sperm parameters (Motrich et al., 2006).

In our study, ROS level and normal morphology were significantly improved after treatment. Parameters such as; was decreased leukocyte count, was increased Sperm count, was increased total motility, was increased class A (rapid progressive), was decreased class D (immotile), was increased Semen volume and was increased acidification of Semen, however, those parameters were not proven statistically. According to Gdoura the mean of Semen volume, Sperm concentration, viability, motility, morphology, and the WBC count were not affected by chlamydial infection (Gdoura et al., 2008). High leukocyte counts in Semen may be an inflammatory marker in the male reproductive system (Aghazarian et al., 2013). The Centers for Disease Control and Prevention recommends azithromycin or doxycycline for the treatment of C.trachomatis infection in which the latter was used, in this study.

An abnormal increase in Semen leukocytes due to was increased ROS levels could lead to Sperm damage (Saleh et al., 2002) (Aitken et al., 2003). In our study, pre-treatment ROS levels were high compared with post-treatment, and ROS was decreased significantly after drug administration and infection removal.

## MATERIALS AND METHODS

### Patient enrolment & ethical statement

The ethical approval for patient enrolment in this study was obtained from the Ethics Committee of Royan institute (IR.ACECR.ROYAN.REC.1394.84). The patients in this study were selected from men consecutively admitted to the Royan Institute in Tehran, Iran. All participants as well as their sexual partners provided written consent letter.

### Inclusion - exclusion criteria, Semen collection and analysis

All the patients were clinically examined and asked for past medical, sexual, and social histories. The study population consisted of men who referred to Royan Institute with Chlamydia trachomatis infection. Eight hundred and forty-eight patients with normal and abnormal Sperm parameters (low Sperm count, pyoSpermia, low Sperm progressive motility, low normal morphology) were included.

Patients with symptoms such as genitourinary tract infections, reproductive system abnormalities, varicocele, testicular tumors, systemic diseases, non-compliance with Spermogram test conditions, and those with a history of antibiotic use in the previous week were not included in our study.

This study was a cross-sectional study, Semen samples were collected into sterile sample cups through self-administered masturbation, after 3–7 days of sexual abstinence. Samples were put in the incubator directly for liquefaction and then manually analyzed by the same person for volume, viscosity, pH, presence of white blood cells (WBCs), Sperm concentration (count per ml and total count), motility (classes A, B, A+B, C, and total), and normal morphology, as indicated by the latest WHO manual for Semen analysis (Organization, 2010). Semen analysis was confirmed using a light microscope equipped with a Computer-Aided Semen Analysis (CASA; Test Sperm2.1, Video test, St. Petersburg, Russia) system. The presence of leukocytes in seminal fluid was detected by peroxidase test. Sperm morphology was detected by staining papanicolaou procedure.

In the first appointment, Semen samples from infertile men were collected in sterile containers and each sample was divided into two parts; the first part for Semen analysis and the next part for Sperm parameters. To evaluate the first part, Sperms were kept into sterile vials in order to perform the Elisa test and PCR at −70 °C until testing was contained. After the centrifugation, plasma samples were analyzed by Elisa test samples and sediment samples were used for DNA extraction. After the infection confirmation by PCR, patients were asked to visit the second visit interval of 3 days from the last ejaculation for Sperm analysis and ROS tests. Antibiotics (every 12 h for two weeks) were also prescribed for them. After the completion of antibiotic usage, If the patients were not resolved from the infection, the treatment continued taking the same antibiotic with the same dose for another week. In order to assess the effect of the empirical antibiotic treatment on Semen parameters ROS levels, as well as clearance from infection, a subsequent Semen sample was taken 30 days after completion of the antibiotic therapy (Pajovic, Radojevic, Vukovic, & Stjepcevic, 2013), by considering the 3–7 days of sexual abstinence.

### Elisa test

To detect specific IgA, antibodies to C. trachomatis by Elisa (Vircell kit IgA: A 1017) a serological assay kit was used and negative controls on each slide were included from the kit. Sample OD was higher (Cut off control: <0.4, 1.5<) in patients who were suspected of being infected or near Cut of (0.4). The DNA extraction and PCR were performed to confirm the presence of infection by Nested PCR.

### Nested PCR

Nested plasmid PCR for C. trachomatis was conducted according to a previously published method on all extracted DNA from Semen. Two pairs of primers were used to detect C. trachomatis as previously described. Products were analyzed by gel electrophoresis in 1.0 % agarose with sybr green staining. DNA of C. trachomatis was extracted from Semen samples using QIAamp mini kit: 51306 according to the manufacturer’s instructions. A negative Control of Extraction was used in the extraction procedure and Internal Control, which served as extraction and amplification control, was directly added to the sample/ lysis mixture during the DNA extraction. Absorbance readings and Genomic detection of Chlamydia trachomatis in Semen samples were conducted by spectrophotometer and Nested PCR, respectively. In brief, dNTP (5 mM) takara company, MgCl_2_ (25 mM) takara company, PCR Buffer (10X) takara company, Taq DNA Polymerase innotrain company, F1/R1،F2/R2 primers (Farayand Danesh co) were used in this study. The contents of Master Mix (25 μl); PCR buffer (10X) 2.5 μl, Mgcl_2_ (25 mM) 1.5μl, dNTPs Mix (10 mM) 0.7 μl, Forward primer (10 pmol/μl) 1.6μl, Reverse primer (10 pmol/μl) 1.6μl, Smar Taq DNA Polymerase (50u/μl) 0.5μl, dd H_2_O 11.6 μl, DNA template 5 μl. PCR Amplification Protocol: Initiation Denaturation for 94 ^°^C for 5 min 1 Cycles, Denaturation 94 ◦C for 30 sec 40 Cycles, Annealing 59 ^◦^C for 30 sec 40 Cycles, Extension 72 ^◦^C for 30 sec 40 Cycles, Final Extension for 72 ^◦^C for 7 min 1 Cycles. Primers: F1 (5’-GGACAAATCGTATCTCGG-3’) R1 (5’-GAAACCAACTCTACGCTG-3’) F2 (5’-ATCCATTGCGTAGATCTCCG-3’) R2 (5’-GCCATGTCTATAGCTAAAGC-3’).

### ROS measurement

Measurement of ROS in Semen specimens was performed according to the WHO laboratory manual for the examination and processing of human Semen (Organization, 2010). Spermatozoa were washed in Krebs–Ringer medium (KRM) and adjusted to 10 ×10^6^ Spermatozoa per ml. Chemiluminescent probes, including luminal, formyl-methionyl-leucyl-phenylalanine (FMLP), and Phorbol 12 myristate 13-acetate (PMA) were utilized to detect extracellular, WBCs and Spermatozoa generated ROS, respectively. Chemiluminescent signals were monitored using a luminometer (SynergyTM H_4_ Hybrid Multi-Mode Microplate Reader, BioTek, USA), and final ROS level was calculated as relative light units Sperm. (RLU/sec/× 10^6^ Sperm).

### Antibiotic treatment and patients’ follow-up

Patients and their sexual partner detected positive for C. trachomatis, were treated with doxycycline (Razak Laboratories, Tehran, Iran), 100 mg orally twice daily for 7 days (Feodorova et al., 2018) (Mishori et al., 2012). To assess the effect of the empirical antibiotic treatment on Semen parameters, ROS levels, as well as clearance from infection, a subsequent Semen sample was taken 30 days after completion of the antibiotic therapy (Pajovic et al., 2013), also considering the 3–7 days of sexual abstinence (Ahmadi et al., 2018). Semen analysis, PCR, ROS assessment, as well as calculation of ROS ratio, were performed on these specimens, as already described for the first samples.

### Statistical method

Non-parametric test Wilcoxon (non-parametric statistical test to compare two dependent groups) was used to analyze the data. The appropriate test plans before and after (a sample on two occasions), non-parametric Wilcoxon test considers the size of the difference between ranking. The P-value <0.05 was considered as statistically significant.

## Abbreviations

CT: Chlamydia trachomatis
ROS: Reactive oxygen species
Elisa: Enzyme-linked immunosorbent assay
PCR: Polymerase chain reaction

## Acknowledgements

This research has been supported by supported by Royan institute for Reproductive Biomedicine.

## Ethics approval

The ethical approval for patient enrolment in this study was obtained from the Ethics Committee of Royan institute (IR.ACECR.ROYAN.REC.1394.84).

## Author contribution

Analyzed the data, evaluated the results, and wrote the paper: RA

## Declarations

### Consent to publish

The following authors have affiliations with organizations with direct or indirect financial interest in the subject matter discussed in the manuscript Authors’ Contributions:

## References

1. Agarwal, A., Sharma, R. K., Nallella, K. P., Thomas Jr, A. J., Alvarez, J. G., & Sikka, S. C. (2006). Reactive oxygen species as an independent marker of male factor Infertility. Fertility and sterility, 86(4), 878–885. https://doi.org/https://doi.org/10.1016/j.fertnstert.2006.02.111

2. Aghaizu, A., Reid, F., Kerry, S., Hay, P. E., Mallinson, H., Jensen, J. S., Kerry, S., Kerry, S., & Oakeshott, P. (2014). Frequency and risk factors for incident and redetected Chlamydia trachomatis infection in sexually active, young, multi-ethnic women: a community based cohort study. Sex Transm Infect, 90(7), 524–528. https://doi.org/http://dx.doi.org/10.1136/sextrans-2014-051607

3. Aghazarian, A., Stancik, I., Pflüger, H., & Lackner, J. (2013). Influence of pathogens and moderate leukocytes on seminal interleukin (IL)-6, IL-8, and Sperm parameters. International urology and nephrology, 45(2), 359–365. https://doi.org/https://doi.org/10.1007/s11255-013-0400-8

4. Ahmadi, M., Mirsalehian, A., Sadighi Gilani, M., Bahador, A., & Afraz, K. (2018). Association of asymptomatic Chlamydia trachomatis infection with male Infertility and the effect of antibiotic therapy in improvement of Semen quality in infected infertile men. Andrologia, 50(4), e12944. https://doi.org/https://doi.org/10.1111/and.12944

5. Aitken, R. J., Baker, M. A., & Sawyer, D. (2003). Oxidative stress in the male germ line and its role in the aetiology of male Infertility and genetic disease. Reproductive biomedicine online, 7(1), 65–70. https://doi.org/https://doi.org/10.1016/S1472-6483(10)61730-0

6. Al-Mously, N., Cross, N. A., Eley, A., & Pacey, A. A. (2009). Real-time polymerase chain reaction shows that density centrifugation does not always remove Chlamydia trachomatis from human Semen. Fertility and sterility, 92(5), 1606–1615. https://doi.org/https://doi.org/10.1016/j.fertnstert.2008.08.128

7. Berktas, M., Aydin, S., Yilmaz, Y., Cecen, K., & Bozkurt, H. (2008). Sperm motility changes after coincubation with various uropathogenic microorganisms: an in vitro experimental study. International urology and nephrology, 40(2), 383. https://doi.org/https://doi.org/10.1007/s11255-007-9289-4

8. Bezold, G., Politch, J. A., Kiviat, N. B., Kuypers, J. M., Wolff, H., & Anderson, D. J. (2007). Prevalence of sexually transmissible pathogens in Semen from asymptomatic male Infertility patients with and without leukocytoSpermia. Fertility and sterility, 87(5), 1087–1097. https://doi.org/https://doi.org/10.1016/j.fertnstert.2006.08.109

9. Bohring, C., & Krause, W. (2003). Immune Infertility: towards a better understanding of Sperm (auto)-immunity: The value of proteomic analysis. Human reproduction, 18(5), 915–924. https://doi.org/https://doi.org/10.1093/humrep/deg207

10. Cunningham, K. A., & Beagley, K. W. (2008). Male genital tract chlamydial infection: implications for pathology and Infertility. Biology of reproduction, 79(2), 180–189. https://doi.org/https://doi.org/10.1095/biolreprod.108.067835

11. de Barbeyrac, B., Papaxanthos-Roche, A., Mathieu, C., Germain, C., Brun, J. L., Gachet, M., Mayer, G., Bébéar, C., Chene, G., & Hocké, C. (2006). Chlamydia trachomatis in subfertile couples undergoing an in vitro fertilization program: a prospective study. European Journal of Obstetrics & Gynecology and Reproductive Biology, 129(1), 46–53. https://doi.org/https://doi.org/10.1016/j.ejogrb.2006.02.014

12. De Francesco, M. A., Negrini, R., Ravizzola, G., Galli, P., & Manca, N. (2011). Bacterial species present in the lower male genital tract: a five-year retrospective study. The European Journal of Contraception & Reproductive Health Care, 16(1), 47–53. https://doi.org/https://doi.org/10.3109/13625187.2010.533219

13. Eley, A., Hosseinzadeh, S., Hakimi, H., Geary, I., & Pacey, A. (2005). Apoptosis of ejaculated human Sperm is induced by co-incubation with Chlamydia trachomatis lipopolysaccharide. Human reproduction, 20(9), 2601–2607. https://doi.org/https://doi.org/10.1093/humrep/dei082

14. Feodorova, V., Sultanakhmedov, E., Saltykov, Y., Zaitsev, S., Utz, S., Corbel, M., Gaydos, C., Quinn, T., & Motin, V. (2018). First detection of chlamydia trachomatis’ Swedish’variant (nvCT) in a Russian couple with Infertility. The open microbiology journal, 12, 343. https://doi.org/10.2174/1874285801812010343

15. Gallegos, G., Ramos, B., Santiso, R., Goyanes, V., Gosálvez, J., & Fernández, J. L. (2008). Sperm DNA fragmentation in infertile men with genitourinary infection by Chlamydia trachomatis and Mycoplasma. Fertility and sterility, 90(2), 328–334. https://doi.org/https://doi.org/10.1016/j.fertnstert.2007.06.035

16. Gdoura, R., Kchaou, W., Ammar-Keskes, L., Chakroun, N., Sellemi, A., Znazen, A., Rebai, T., & Hammami, A. (2008). Assessment of Chlamydia trachomatis, Ureaplasma urealyticum, Ureaplasma parvum, Mycoplasma hominis, and Mycoplasma genitalium in Semen and first void urine specimens of asymptomatic male partners of infertile couples. Journal of andrology, 29(2), 198–206. https://doi.org/https://doi.org/10.2164/jandrol.107.003566

17. Hislop, J., Quayyum, Z., Flett, G., Boachie, C., Fraser, C., & Mowatt, G. (2010). Systematic review of the clinical effectiveness and cost-effectiveness of rapid point-of-care tests for the detection of genital chlamydia infection in women and men. Health Technology Assessment. https://doi.org/10.3310/hta14290

18. Hosseinzadeh, S., Eley, A., & Pacey, A. A. (2004). Semen quality of men with asymptomatic chlamydial infection. Journal of andrology, 25(1), 104–109. https://doi.org/https://doi.org/10.1002/j.1939-4640.2004.tb02764.x

19. Idahl, A., Abramsson, L., Kumlin, U., Liljeqvist, J., & Olofsson, J. (2007). Male serum Chlamydia trachomatis IgA and IgG, but not heat shock protein 60 IgG, correlates with negatively affected Semen characteristics and lower pregnancy rates in the infertile couple. International Journal of Andrology, 30(2), 99–107. https://doi.org/https://doi.org/10.1111/j.1365-2605.2006.00718.x

20. Kalantar, S. M., Kazemi, M. J., Sheikhha, M. H., Aflatoonian, A., & Kafilzadeh, F. (2007). Detection of Chlamydia trachomatis infection in female partners of infertile couples. Int J Fertil Steril, 1(2). https://doi.org/http://ijfs.ir/journal/article/abstract/2298.

21. Kalayoglu, M. V., & Byrne, G. I. (2006). The genus Chlamydia-medical. The prokaryotes, 7, 741–754. https://doi.org/https://doi.org/10.1007/0-387-30747-8_30

22. Khosrowbeygi, A., & Zarghami, N. (2007). Levels of oxidative stress biomarkers in seminal plasma and their relationship with seminal parameters. BMC clinical pathology, 7(1), 6. https://doi.org/https://doi.org/10.1186/1472-6890-7-6

23. Mahfouz, R. Z., du Plessis, S. S., Aziz, N., Sharma, R., Sabanegh, E., & Agarwal, A. (2010). Sperm viability, apoptosis, and intracellular reactive oxygen species levels in human Spermatozoa before and after induction of oxidative stress. Fertility and sterility, 93(3), 814–821. https://doi.org/https://doi.org/10.1016/j.fertnstert.2008.10.068

24. Mazzoli, S., Cai, T., Addonisio, P., Bechi, A., Mondaini, N., & Bartoletti, R. (2010). Chlamydia trachomatis infection is related to poor Semen quality in young prostatitis patients. European urology, 57(4), 708–714. https://doi.org/https://doi.org/10.1016/j.eururo.2009.05.015

25. Mishori, R., McClaskey, E. L., & WinklerPrins, V. (2012). Chlamydia trachomatis infections: screening, diagnosis, and management. American family physician, 86(12), 1127–1132. https://doi.org/https://www.aafp.org/afp/2012/1215/p1127.html

26. Motrich, R. D., Cuffini, C., Oberti, J. P. M., Maccioni, M., & Rivero, V. E. (2006). Chlamydia trachomatis occurrence and its impact on Sperm quality in chronic prostatitis patients. Journal of Infection, 53(3), 175–183. https://doi.org/https://doi.org/10.1016/j.jinf.2005.11.007

27. Muriel, L., Garrido, N., Fernández, J. L., Remohí, J., Pellicer, A., de los Santos, M. J., & Meseguer, M. (2006). Value of the Sperm deoxyribonucleic acid fragmentation level, as measured by the Sperm chromatin dispersion test, in the outcome of in vitro fertilization and intracytoplasmic Sperm injection. Fertility and sterility, 85(2), 371–383. https://doi.org/https://doi.org/10.1016/j.fertnstert.2005.07.1327

28. Oehninger, S. (2000). Clinical and laboratory management of male Infertility: an opinion on its current status. Journal of andrology, 21(6), 814–821. https://doi.org/https://doi.org/10.1002/j.1939-4640.2000.tb03411.x

29. Organization, W. H. (2010). WHO laboratory manual for the examination and processing of human Semen. https://doi.org/https://apps.who.int/iris/bitstream/handle/10665/44261/9789241547789_eng.pdf;jsessionid=E1A703A7F0185327697DBF52383B0D01?sequence=1

30. Pajovic, B., Radojevic, N., Vukovic, M., & Stjepcevic, A. (2013). Semen analysis before and after antibiotic treatment of asymptomatic chlamydia-and ureaplasma-related pyoSpermia. Andrologia, 45(4), 266–271. https://doi.org/https://doi.org/10.1111/and.12004

31. Peeling, R., & Embree, J. (2005). Screening for sexually transmitted infection pathogens in Semen samples. Canadian Journal of Infectious Diseases and Medical Microbiology, 16(2), 73–76. https://doi.org/https://doi.org/10.1155/2005/958374

32. Prabha, V., Sandhu, R., Kaur, S., Kaur, K., Sarwal, A., Mavuduru, R. S., & Singh, S. K. (2010). Mechanism of Sperm immobilization by Escherichia coli. Advances in urology, 2010. https://doi.org/https://doi.org/10.1155/2010/240268

33. Saleh, R. A., Agarwal, A., Kandirali, E., Sharma, R. K., Thomas Jr, A. J., Nada, E. A., Evenson, D. P., & Alvarez, J. G. (2002). LeukocytoSpermia is associated with was increased reactive oxygen species production by human Spermatozoa. Fertility and sterility, 78(6), 1215–1224. https://doi.org/https://doi.org/10.1016/S0015-0282(02)04237-1

34. Schillinger, J. A., Dunne, E. F., Chapin, J. B., Ellen, J. M., Gaydos, C. A., Willard, N. J., Kent, C. K., Marrazzo, J. M., Klausner, J. D., & Rietmeijer, C. A. (2005). Prevalence of Chlamydia trachomatis infection among men screened in 4 US cities. Sexually transmitted diseases, 32(2), 74–77. https://doi.org/doi:10.1097/01.olq.0000149670.11953.ca

35. Shiva, M., Gautam, A. K., Verma, Y., Shivgotra, V., Doshi, H., & Kumar, S. (2011). Association between Sperm quality, oxidative stress, and seminal antioxidant activity. Clinical biochemistry, 44(4), 319–324. https://doi.org/https://doi.org/10.1016/j.clinbiochem.2010.11.009

36. Stamm, W. E., Batteiger, B. E., Mccormack, W. M., Totten, P. A., Sternlicht, A., Kivel, N. M., & Group, R. S. (2007). A randomized, double-blind study comparing single-dose rifalazil with single-dose azithromycin for the empirical treatment of nongonococcal urethritis in men. Sexually transmitted diseases, 34(8), 545–552. https://doi.org/doi:10.1097/01.olq.0000253348.44308.8c

37. Tavilani, H., Doosti, M., & Saeidi, H. (2005). Malondialdehyde levels in Sperm and seminal plasma of asthenozooSpermic and its relationship with Semen parameters. Clinica chimica acta, 356(1-2), 199–203. https://doi.org/https://doi.org/10.1016/j.cccn.2005.01.017

38. Tirado, E. E., Sharma, R., Sawyer, D. E., Awasthi, Y. C., & Brown, D. B. (2003). Effects of oxidative stress on human Sperm activation. Fertility and sterility, 80, 240. https://doi.org/DOI:https://doi.org/10.1016/S0015-0282(03)01581-4

39. Watson, E. J., Templeton, A., Russell, I., Paavonen, J., Mardh, P.-A., Stary, A., & Pederson, B. S. (2002). The accuracy and efficacy of screening tests for Chlamydia trachomatis: a systematic review. Journal of medical microbiology, 51(12), 1021–1031. https://doi.org/https://doi.org/10.1099/0022-1317-51-12-1021

40. Zhang, Z., Zhang, H., Dong, Y., Han, R., Dai, R., & Liu, R. (2011). Ureaplasma urealyticum in male Infertility in Jilin Province, North-east China, and its relationship with Sperm morphology. Journal of International Medical Research, 39(1), 33–40. https://doi.org/https://doi.org/10.1177/147323001103900104

